# Temporal dynamics of ensemble statistics calculation using a neural network model

**DOI:** 10.1101/2020.04.20.051151

**Authors:** Rakesh Sengupta

## Abstract

Computing summary or ensemble statistics of a visual scene is often automatic and a hard necessity for stable perceptual life of a cognitive agent. Although computationally the process should be as simple as applying a filter as it were to a perceived scene, the issue of mechanism of summary statistics is complicated by the fact that we can seamlessly switch from summarizing to individuation while computing the ensemble averages across multiple reference frames. In the current work we have investigated the possibility of a neural network that can also switch between individuation and summarization. We have chosen a computational model previously used for enumeration/individuation (Sengupta et al, 2014) in order to show possibility of extracting summary statistics using two different measures from the network. The results also shed a light on possible temporal dynamics of ensemble perception.

## Introduction

Visual system works with two mechanisms, one with detailed individuation of few items within attentional focus, and computation of average statistics of scenes outside attentional focus. Both combine to give a stable perceptual world for human beings (Corbett and Melcher, 2013, 2014). There are advantages to representing ensemble averages of features for similar and redundant items in line with principles of cognitive economy. That the averaging task is undoubtedly less demanding compared to individuation, is supported by the fact that membership judgement tasks are found to be three times more difficult than perceptual averaging task (Ariely, 2001). As such, average properties of sets of similar objects can be represented without retaining information about the individual items comprising the sets, even when the individual elements cannot be consciously perceived (Alvarez and Oliva, 2008; Corbett and Oriet, 2011). This averaging perceptual activity, at least in part, seems to be part of default sensory activity. For instance, Oriet and Brand (2013) have shown that even under explicit instructions observers could not disengage from perceptual averaging activity. Corbett and Melcher (2014) have also shown how the summary statistics for mean size adaptation operates across multiple reference frames, retinotopic, spatiotopic and hemispheric. Brady and Alvarez (2011) have also pointed out that even representations in visual working memory are biased through ensemble statistics. There is also evidence that global ensemble statistics, rather than object based or non-spatial global cues, are critical to scene perception (Brady et al., 2017).

In terms of mechanism, it has been suggested that the joint statistics for peripheral stimuli is collected from differential response from neural assemblies sensitive to changes in position, phase, orientation and scale (Rosenholtz et al., 2012). However, there exists some disagreement regarding the temporal dynamics of the cognitive process. Whereas Chong and Treisman (2003) propose a faster automatic process (within ∼50 ms), Whiting and Oriet (2011) on the other hand suggest a slower timeline (around ∼200 ms). In the current work we investigated this dichotomy through a computational model developed for individuation (see Sengupta et al., 2014 for details). We argue that since our perceptual system handles the transition between ensemble averaging and individuation quite seamlessly in the course of everyday perception, it holds merit to investigate neural mechanisms that may serve purpose of both of these cognitive activities. In the following section we will show how such a model dynamics can be built and the consequences towards mechanisms of summary statistics.

## Results

In our previous work (Sengupta et al., 2014), we proposed a recurrent on-center off-surround architecture based on fixed self-excitation (*α*) and lateral inhibition (*β*) in a fully connected network. The temporal dynamics of the activation of *i*-th node is given by,

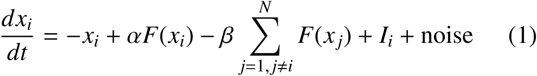

The activation function *F*(*x*) is given by

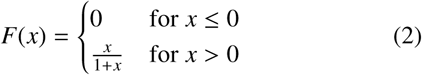

Input to the *i*-th node is given by *I*_*i*_, which is has value if a particular node is given a input at a particular time and 0 otherwise. Noise is sampled at every time step from a Gaussian distribution with mean 0 and standard deviation 0.05. In the present work, the input to the network consisted of inputs sampled for the number of required inputs (set size parameter in Table 1) from a Gaussian distribution with mean 0.3 and and standard deviation of 0.1. The stimulus presentation time was varied between 60 and 220 ms. See Table 1 for simulation parameters^1^. Schematics of the network are given in Fig. 1. The model dynamics was computed using Euler method with step size 0.01.

**Table 1.**
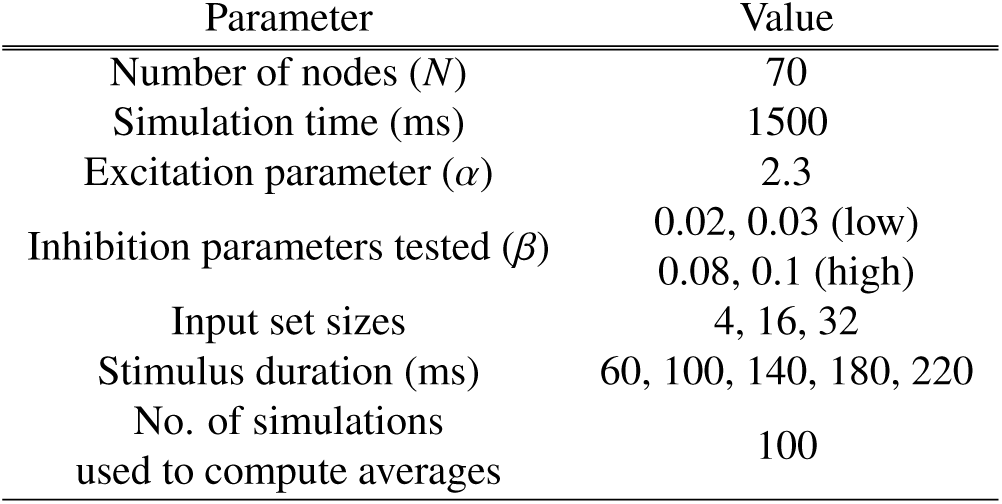
Model parameters

**Figure 1.**
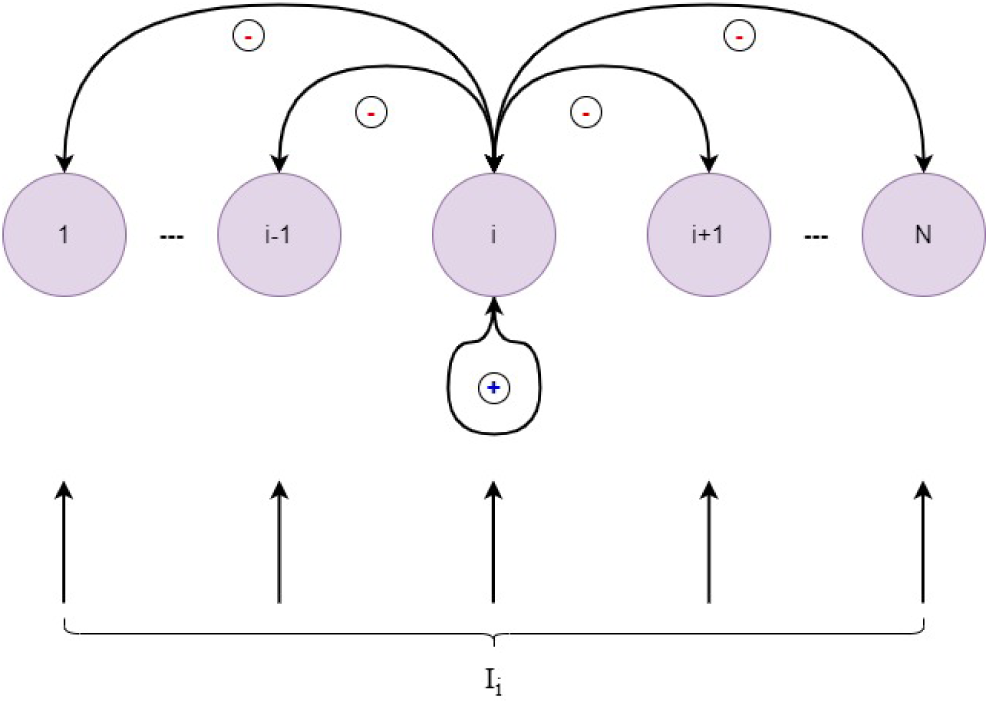
The computational model used for summary statistics calculation.

The ensemble statistics related results were calculated using two measures. We stored input and output (network output at steady state/ end of simulation) of the network as vector strings mapped to binary 0/1 values (input or output values above 0.05 was stored as 1, otherwise 0). First measure of ensemble statistics was Euclidean distance between the input and output strings, and the other measure was the *r*^2^ value computed between the two strings. The rational behind the choice was that the computing more accurate summary statistics. For each set seize and each inhibition parameter, the simulation was run 100 times to calculate average euclidean distance and *r*^2^.

The results shown in Fig. 2 reaffirms previously known behavior for such networks - such as high inhibition is better for small set sizes and low inhibition is better for enumerating larger set sizes. Thus it comes as no surprise that euclidean distance measure for set size 4 is much lower at higher inhibition than at low inhibition. Similar results are also seen for correlation measure. For larger set sizes 16 and 32, we can see that euclidean distance decreases with increasing presentation time with sharper decline for set size 16 (see Fig. 2a). A similar trend can be seen with the correlation calculation as well (see Fig. 2b).

**Figure 2.**
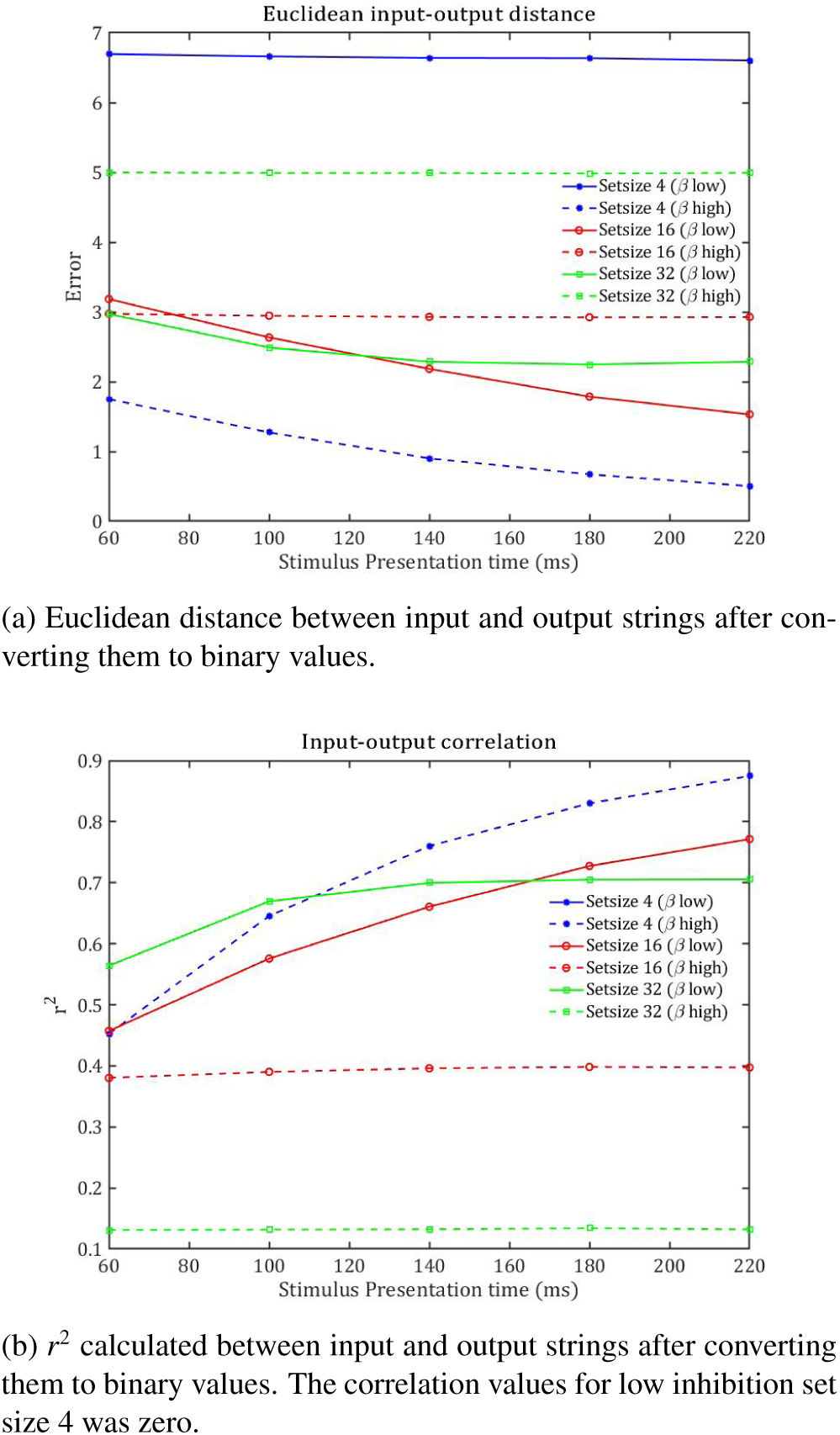
Two measures of ensemble statistics calculated from the model. For low inhibition result, the average of *β* = 0.02 and 0.03 was used. We used average of *β* = 0.08 and 0.1 to compute high inhibition result.

## Discussion

In the current work, we have used the same neural network model used in our previous work on enumeration (Sengupta et al., 2014, 2017), in order to understand possible strategies for computing summary statistics in the brain. While it is understandable that simple feed-forward filters make more sense computationally when we try to digitally summarize images, the crucial fact remains that our perceptual system is flexible enough to switch seamlessly between individuation and computing summary statistics. In order for such switch to be possible, a certain topographical similitude between input and output to the neural substrate is necessary. Originally we had devised the model (see Fig. 1) to emulate topographical functions of lateral intraparietal cortex (LIP) seen during enumeration tasks (Knops et al., 2014). Detailed precise enumeration requires individuation and thus high inhibition, however less precise estimation of numerosity requires lower inhibition. We compared the relation between the input and output vector to the network at low inhibition range with two different measures explained in previous section and found that the network dynamics conforms to the behavior suggested by Whiting and Oriet (2011). For larger set sizes the *r*^2^ value crosses higher values like 0.7 only at presentation times greater than 100 ms. A more complete planned model for computing summary statistics requires the implementation within a larger scope of an architecture along with the individuation mechanism with explicitly coded transition between them. We intend to pursue such a model in our future work.

## Footnotes

1 Details regarding the parameter choices are given in Appendix of Sengupta et al. (2014).

## References

Alvarez, G. A. and Oliva, A. (2008). The representation of simple ensemble visual features outside the focus of attention. Psychological Science, 19:392–398.

Ariely, D. (2001). Seeing sets: representation by statistical properties. Psychol Sci, 12(2):157–162.

Brady, T. F. and Alvarez, G. A. (2011). Hierarchical encoding in visual working memory: Ensemble statistics bias memory for individual items. Psychological science, 22(3):384–392.

Brady, T. F., Shafer-Skelton, A., and Alvarez, G. A. (2017). Global ensemble texture representations are critical to rapid scene perception. Journal of Experimental Psychology: Human Perception and Performance, 43(6):1160.

Chong, S. C. and Treisman, A. (2003). Representation of statistical properties. 43:393–404.

Corbett, J. and Melcher, D. (2013). Summary statistics support spatiotemporal stability. Journal of Vision, 13:1043–1043.

Corbett, J. E. and Melcher, D. (2014). Characterizing ensemble statistics: mean size is represented across multiple frames of reference. Atten Percept Psychophys, 76:746–758.

Corbett, J. E. and Oriet, C. (2011). The whole is indeed more than the sum of its parts: perceptual averaging in the absence of individual item representation. Acta Psychol (Amst), 138(2):289–301.

Knops, A., Piazza, M., Sengupta, R., Eger, E., and Melcher, D. (2014). A shared, flexible neural map architecture reflects capacity limits in both visual short-term memory and enumeration. J Neurosci, 34(30):9857–9866.

Oriet, C. and Brand, J. (2013). Size averaging of irrelevant stimuli cannot be prevented. Vision Res, 79:8–16.

Rosenholtz, R., Huang, J., Raj, A., Balas, B. J., and Ilie, L. (2012). A summary statistic representation in peripheral vision explains visual search. Journal of Vision, 12:1–17.

Sengupta, R., Bapiraju, S., and Melcher, D. (2017). Big and small numbers: Empirical support for a single, flexible mechanism for numerosity perception. 79:253–266.

Sengupta, R., Surampudi, B. R., and Melcher, D. (2014). A visual sense of number emerges from the dynamics of a recurrent on-center off-surround neural network. 1582:114–124.

Whiting, B. F. and Oriet, C. (2011). Rapid averaging? not so fast! 18:484–489.

